# Exploring Auditory Category Distinctions in Perception and Imagery

**DOI:** 10.1101/2025.07.21.665330

**Authors:** Julia M. Leeman, Evan Hare, Ricardo Morales-Torres, Pooja Kabber, Joseph Zhang, Kristi Van Meter, Tobias Overath

**Author notes:** **Corresponding Authors:** Julia Leeman, Evan Hare. These authors contributed equally to this work (Julia M. Leeman, Evan Hare).

## Abstract

Imagery, the ability to generate perceptual experiences in the absence of external stimuli, is used every day when remembering a past event or imagining a novel situation. While most previous research on imagery has focused on the visual domain, the present study presents an investigation of auditory imagery. The perception of different categories of sound has been shown to evoke different neural responses. Further, the neural processes underlying auditory imagery and perception have been shown to be similar. Therefore, we hypothesized that auditory imagery would rely on similar categorical processing. Participants learned shape-sound associations and were then asked to imagine the matching sound when presented with the associated shape. We chose two example stimuli from two maximally different sound categories — human speech sounds and nonhuman environmental sounds — to investigate our hypothesis. Electroencephalography (EEG) data were recorded while participants listened to and imagined the sounds. The mean voltage in the P2 event-related potential time window (180-280 ms) was significantly larger for perception of the speech sounds than the environmental sounds, and the late positive event-related potential complex (LPC, 350-500 ms) associated with imagery was significantly smaller for imagery of the speech sounds than the environmental sounds. This suggests that, as in perception, the neural processing of imagined sounds is categorical.

## 1 Introduction

You’ve been on a long trip abroad and, as you walk up to your door, you imagine your dog barking as it rushes towards you and your loved ones shouting, “Welcome home!” This experience of sound in the absence of external stimuli is called auditory imagery. Auditory imagery has been shown to encode many of the same properties as auditory perception, including pitch and loudness (Wu et al., 2011). However, it is unknown whether auditory imagery and perception rely on similar categorical distinctions. The present study aims to answer this question by recording electroencephalography (EEG) data while participants imagined sounds from different categories.

To respond behaviorally to sounds, we must classify them into different categories. For example, to understand words, it is necessary to first recognize the sound as human speech. Categorical processing is present throughout the auditory system, with core auditory regions responding to more simple categorical features, such as frequency contours, and regions further along the auditory ventral pathway responding to more complex auditory categories (Tsunada & Cohen, 2014). One of the main categorical distinctions found in human auditory perception is differential processing of the human voice. Areas in the superior temporal sulcus bilaterally are activated significantly more by human vocal sounds than acoustically matched non-vocal sounds (Belin et al., 2000). Similarly, as compared to fundamental-frequency-matched instrumental tones, sung tones elicited a greater late positive event-related potential (ERP), termed the “voice-specific response” (Levy et al., 2001). Charest et al. (2009) suggested that voice selectivity begins earlier, with the “fronto-temporal positivity to voices” early latency ERP in response to the human voice (human speech sounds and vocalizations) having a greater amplitude than responses to other sounds (bird songs, natural sounds, mechanical sounds, and instruments). Despite extensive evidence for category-specific processing in auditory perception, this has not been well-studied in the domain of imagery aside from a recent study showing the effect of emotional content on auditory imagery (Proverbio et al., 2023).

There are many similarities between perception and imagery. For example, auditory imagery has been demonstrated for various perceptual properties of sound, such as timbre (Hubbard, 2010; Tużnik et al., 2018), loudness (Hubbard, 2010; Tian et al., 2018; Wu et al., 2011), pitch (Halpern, 2015; Hubbard, 2010; Wu et al., 2011) and melody (Zatorre et al., 2010). Additionally, imagery and perception recruit partly overlapping neural circuits (Dijkstra et al., 2017, 2019; Gelding et al., 2019; McNorgan, 2012; Regev et al., 2021). A recent study (Ding et al., 2019) showed that auditory perception and imagery share similar neural pathways, but follow distinct time courses: while auditory perception relied on bottom-up activation of a pathway from the temporal gyrus to more frontal regions, auditory imagery relied on top-down activation of a pathway from the inferior frontal gyrus to the temporal gyrus. These distinct time courses are reflected in the ERP response. Proverbio et al. (2023) suggested that a late-positive complex (LPC) evoked by imagery reflects categorical differences found in the P2 elicited by auditory perception. However, categorical processing in this study was related to the emotional content of the stimuli (Proverbio et al., 2023). We seek to determine whether imagery distinguishes between auditory semantic categories.

Here, we asked whether the neural correlates of auditory imagery, as measured with EEG, could be distinguished based on sound category. We chose examples of two maximally different sound categories — speech sounds and environmental sounds — to answer this question. We trained participants to associate specific shapes with the sounds, asking them to imagine the matching sound when visually presented with the associated shape. We compared neural responses to the speech and environmental sounds for both perception and imagery. We hypothesized that categorical processing would be evident in the P2 for auditory perception and the LPC for imagery.

## 2 Methods

### 2.1 Participants

Thirty-nine participants (age range: 18-43, mean age: 20.67, 15 males) were recruited for EEG data collection. Nine were rejected for poor behavior or overall poor EEG data quality, leaving thirty remaining participants (mean age: 22.03, 13 males) that were included in this study. All participants gave written informed consent prior to the study, in compliance with the protocols approved by the Duke University Institutional Review Board. Participants were compensated either monetarily or with class credit.

### 2.2 Stimuli

The visual stimuli consisted of six simple geometric shapes: square, circle, star, diamond, half-circle, and triangle, presented on an LCD monitor. Four auditory stimuli were chosen from a list of six stimuli due to a similar vividness, difficulty, and accuracy of auditory imagery (see Appendix). All analyses will focus on two different categories of sound: speech and environmental sounds. There were two exemplars of each sound category: the English words “dig” and “cut”, as well as a single frog vocalization and an isolated car horn. The “dig” and “cut” stimuli were recorded speech from an adult English-speaking male and matched in pitch (∼130 Hz). The two environmental sounds were royalty free sound effects collected from the internet and were matched in average pitch and intensity to the speech sounds. All sounds were 700 ms in duration and were presented at ∼70 dB through Etymotic insert earphones using an RME Fireface UC external audio card at a sampling rate of 44.1 kHz. All visual and auditory stimuli were presented using Psychtoolbox-3 (Brainard, 1997) in MATLAB 2018b (The MathWorks Inc., 2018) on a Mac computer.

### 2.3 Procedure

#### 2.3.1 Validation of Stimuli

We recorded both self-reported auditory imagery experiences as behavioral measures and neural responses measured using EEG. First, participants listened to the auditory stimuli for this study and were asked to briefly describe the stimuli and rate the sounds based on how speechlike they were on a scale from 1 (definitely not speech) to 5 (definitely speech). Next, they were asked to try to imagine the sounds and rate them based on how vivid their mental reproductions of the sounds were, how difficult they were to imagine, and how accurate their mental reproductions of the sounds were.

#### 2.3.2 Subjective Vividness of Auditory Imagery

Using the Bucknell Auditory Imagery Vividness Subscale (BAIS-V), a set of 14 questions addressing musical, verbal, and environmental internal auditory experience, we next collected subjective scores of auditory imagery vividness (Halpern, 2015). The BAIS-V subscale focused on perceived similarity of auditory images to auditory perception. For each auditory image on the BAIS-V, participants were asked to rank vividness on a Likert scale from 1 (no image present at all) to 7 (as vivid as the actual sound).

#### 2.3.3 Training

Data collection was divided into three blocks of training and testing, each pairing two shapes with two sounds. During training, participants sat in a sound-attenuated room and conducted a visual-auditory association task, where shapes were presented and followed 300 ms later by their paired sounds. A fixation cross with a jitter of 600-800 ms followed each visual-auditory presentation. Each trial presented both shape-sound pairs twice over 4 s. Presentation order and associations were counterbalanced across participants. Every six trials, a validation phase occurred: participants saw one shape followed by two sounds in random order and indicated which matched. This phase included 10 trials (5 per shape). Participants who reached ≥90% accuracy advanced to testing; others repeated training.

#### 2.3.4 Testing

During the testing phase, EEG data were recorded while participants were presented with one of the two shapes and asked to imagine their associated sound from their training phase (see Figure 1). Each trial consisted of the presentation of a shape for 2 s, during which the participant was supposed to imagine the associated sound. Then, the participants were presented with one of the two sounds that could either be a match or mismatch with the previous shape (50% chance of match). Participants were then asked to respond using a keyboard whether the sound they heard matched the sound they had imagined. They were immediately given feedback on the screen as to whether their response was correct or incorrect, and then they proceeded to the next trial.

**Figure 1.**
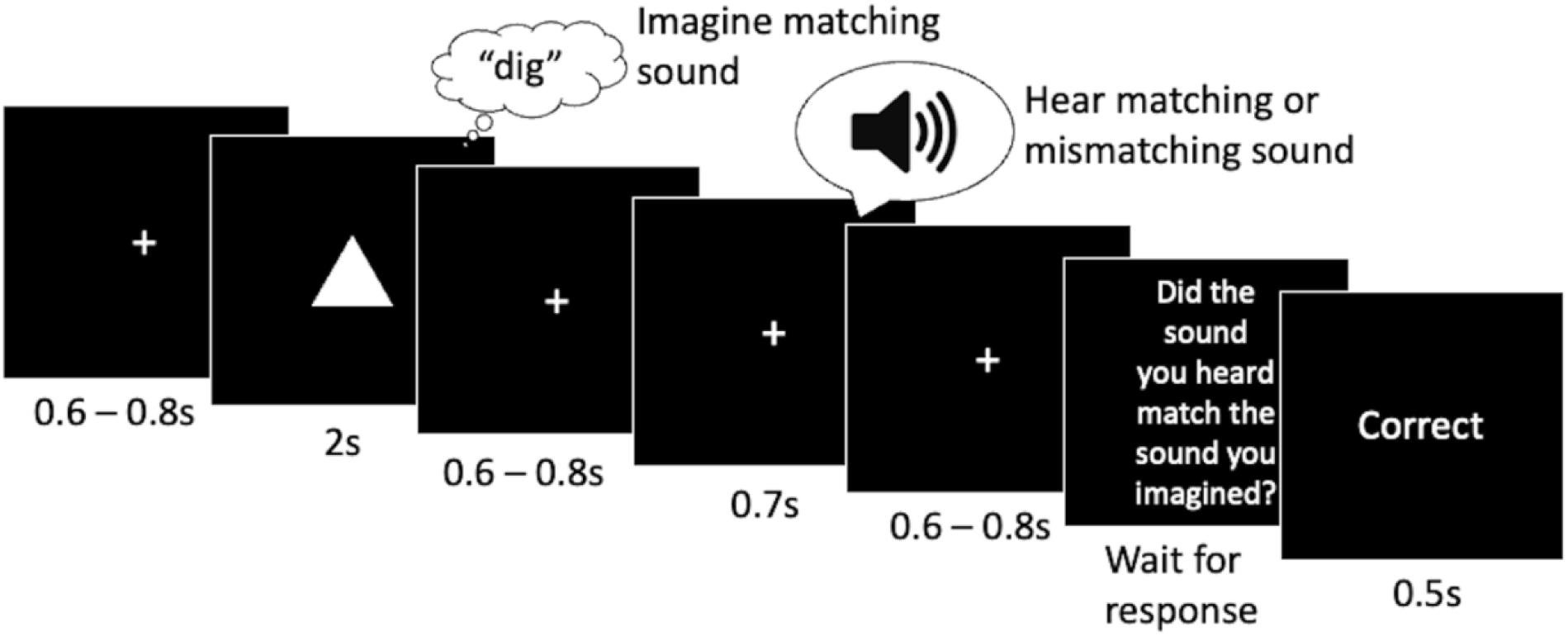
Visual representation of an example trial. Participants were presented a shape and asked to imagine the matching sound. Then, they heard a sound that was either a match or mismatch (50% chance) and were asked to respond whether that sound matched the sound they imagined. Participants were provided feedback.

The test phase consisted of 107 trials (three practice trials, and 104 test trials), each around 6.3 s long, for a total of 11 – 12 mins for each test block. The order of stimulus presentation was randomized, controlling for no greater than three consecutive repetitions of the same shape and no more than three matches or mismatches in a row. Participants completed both training and testing three times with three different image-sound pairs. Each stimulus was presented a total of 52 times (104 per category).

### 2.4 EEG Data Acquisition

EEG was recorded on a 64-channel active electrode system (Brain Vision ActiChamp, Brain Products) using a customized, extended coverage, elastic electrode cap (EASYCAP, Herrsching, Germany). This cap provides extended coverage of the head from just above the eyebrows to below the inion posteriorly and has electrodes that are equally spaced across the cap. Two fronto-lateral electrodes track horizontal eye movements, while an additional external electrode just underneath the left eye tracks vertical eye movements. EEG data were recorded at a 1000 Hz sampling rate and referenced to the right mastoid. Electrode impedances were maintained below 15 kΩ where possible.

### 2.5 Analysis of EEG Data

Data were preprocessed using EEGLAB (Delorme & Makeig, 2004) and custom-written scripts in MATLAB 2021b (The MathWorks Inc., 2021). Data up until 2 seconds before the first trial of each session and as of 2 seconds after the last trial were removed. The data were re-referenced to the average of the left and right mastoids and bandpass filtered between 0.1-50 Hz. Electrodes that exhibited high amounts of noise resulting from poor contact were interpolated using data from nearby electrodes (no more than 2 per participant). Independent component analysis was used to remove non-neural artifacts in the data, including eye blinks, eye movements, and muscle tension. The data were epoched between [-600 2000] ms with respect to the onset of the shape (auditory imagery) and [-600 1300] ms with respect to sound onset (auditory perception). Epochs were baseline corrected between [-200 0] ms with respect to the onset of the stimulus. Trials in which participants reported imagining the sound that did not match with the presented shape were excluded from analysis. In addition, trials that contained artifacts based on visual inspection after ICA were rejected. Rejection never exceeded 10 trials for a single stimulus for a single participant.

ERP data were analyzed using FieldTrip (Oostenveld et al., 2011). Statistical tests run in R included repeated measures analyses of variance (ANOVA) and paired t-tests. The Greenhouse-Geisser correction was used to adjust for violation of the sphericity assumption. Benjamini & Hochberg’s method was used to control the false discovery rate for multiple comparisons. The ERP analyses focused on the amplitude in the N1 (50-150 ms), P2 (180-280 ms) and LPC (350-500 ms) post-stimulus time periods, as in Stavropoulos & Carver (2016) and Wu et al. (2011). The analysis also focused on six predefined regions of interest (ROI) based on Wu et al. (2011): these regions consisted of an anterior-left (F3, FC3, C3), anterior-middle (Fz, FCz, Cz), anterior-right (F4, FC4, C4), posterior-left (CP3, P3, PO3), posterior-middle (CPz, Pz, POz), and posterior-right (CP4, P4, PO4) ROI.

After conducting our hypothesis driven ERP analysis, we conducted a classification analysis employing EEGNet (Lawhern et al., 2018), a convolutional neural network designed to extract spatial and temporal features of the neural time series. We employed this data-driven approach to interrogate whether the imagery of the speech vs. the environmental sounds would elicit distinct patterns of neural activity. To perform this analysis, we trained EEGNet to classify neural responses as evoked by imagery of the speech or environmental sounds. To calculate the classification accuracy, we employed a 10-fold cross-validation approach: training was performed on 90% of the data, while the remaining 10% was used to test whether the learned weights could correctly classify the data into speech or environmental sounds. The Python package DeepExplain (Ancona et al., 2018) was employed to reveal which temporal and spatial regions of the neural time-series were more informative to the classifier accuracy. This package allows users to generate a visual representation of the weights, across electrodes and time-points, that EEGNet used during the classification between the speech and environmental sounds.

Finally, to test whether the classification accuracy was above chance, we employed a permutation approach in which we shuffled the labels (speech vs. environmental) of the trials and performed the 10-fold cross-validation. Then, to construct a distribution of results reflecting a null hypothesis (i.e., a distribution that reflects no difference between conditions), we repeated this process 400 times. We then determined the percentile of our real classification accuracy within this null distribution to assess statistical significance.

## 3 Results

### 3.1 BAIS ratings and Behavior

First, we investigated participants’ scores on the BAIS-V scale. The mean rating across all questions and all participants was 5.083 (range: 3.357 – 6.714, standard deviation: 0.936). Since a rating of 4 represented a fairly vivid image, this suggests that all participants could imagine different kinds of sound. Next, we investigated participants’ behavioral accuracy on the imagery task. Participants responded with a mean accuracy of 97.186% (range: 86.916-100%, standard deviation: 2.917%) to the question “Did the sound you heard match the sound you imagined?” on the test session. This high level of accuracy suggests that participants imagined the sound that matched the shape presented.

### 3.2 Event-Related Potentials

#### 3.2.1 Perception

First, we sought to validate categorical differences in perception found in previous literature (Charest et al., 2009). Grand average ERPs evoked by perception of the speech and environmental sounds are displayed in Figure 2. To investigate the N1, we ran a repeated measures ANOVA with stimulus category (speech vs. environmental sounds), anterior-posterior electrode position, and laterality (left, middle, right) as factors and mean voltage across the N1 time window (50-150 ms) as the dependent variable. This revealed main effects of stimulus (F_1, 29_ = 12.451, p < .01) and laterality (F_2, 58_ = 6.884, p < .01), while both stimulus and anterior-posterior position (F_1, 29_ = 74.832, p < .0001) and stimulus and laterality (F_2, 58_ = 6.348, p < .01) showed an interaction. As seen in Figure 2, the environmental sounds elicited a more negative N1 than speech sounds, especially in the anterior ROIs. Paired t-tests were conducted to inspect the interaction effects. These tests revealed that the mean voltage difference between the speech and environmental sounds was greatest in anterior regions (anterior – posterior: t_179_ = 10, p < 0.001) and in the middle of the scalp (middle – left: t_119_ = 2.7, p < .05, middle – right: t_119_ = 4.3, p < .001).

**Figure 2.**
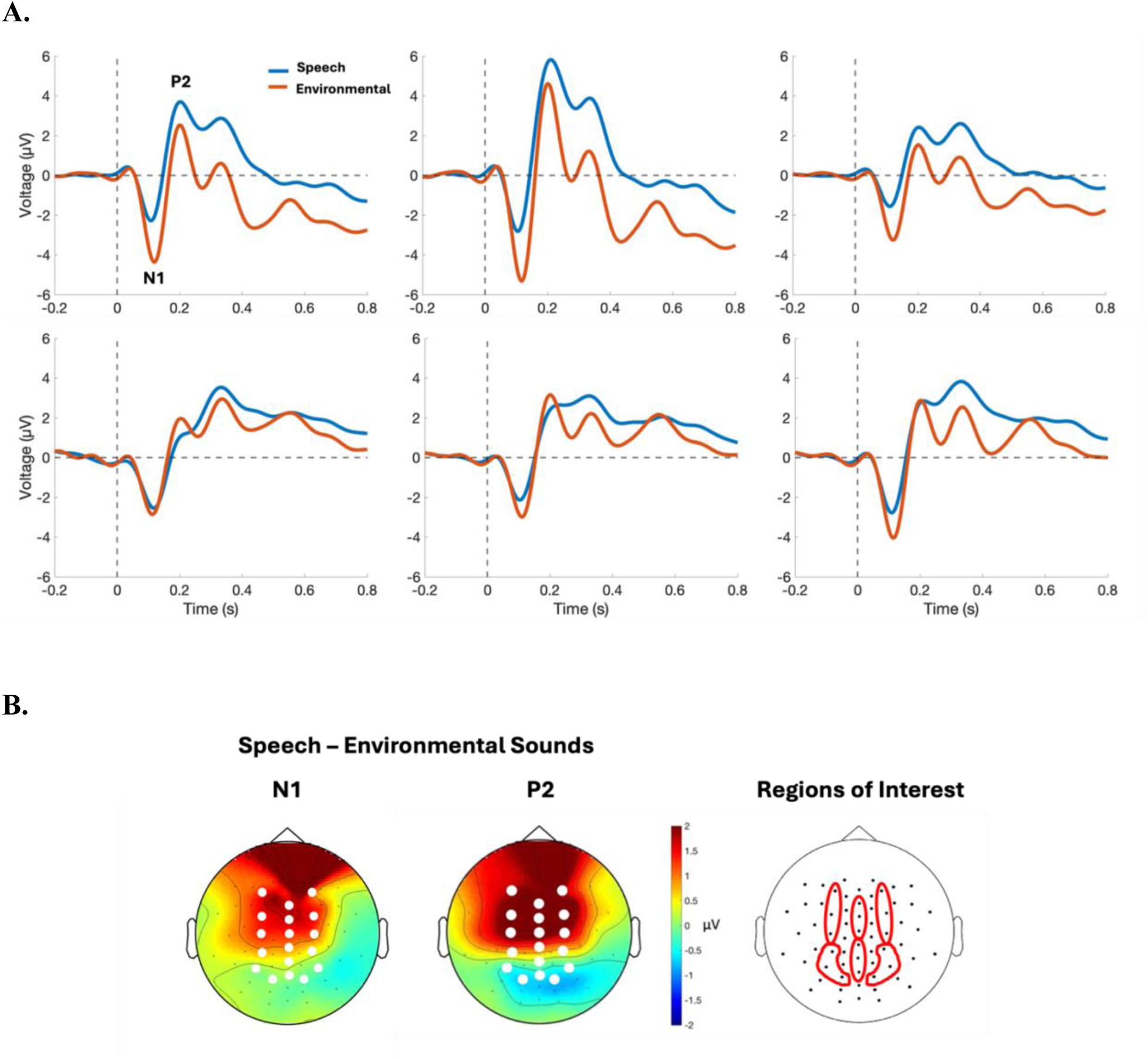
A. Grand average ERPs evoked by perception of the speech and environmental sounds. ERPs are displayed from each ROI. B. The topographic plots represent the response to speech sounds minus the response to environmental sounds in the time windows of interest (N1 and P2). All 6 ROIs visually represented with red lines.

Next, we investigated the P2. We conducted a repeated measures ANOVA on the perception epochs with stimulus category (speech vs. environmental sounds), electrode position (anterior, posterior), and laterality (left, middle, right) as factors and mean voltage across the P2 time window (180-280 ms) as the dependent variable. This revealed main effects of stimulus (F_1, 29_ = 10.429, p < .01) and laterality (F_1.669, 48.401_ = 59.589, p < .001) and interactions between stimulus category and anterior-posterior position (F_1, 29_ = 46.715, p < .001) and stimulus category and laterality (F_2, 58_ = 7.942, p < .001). As seen in Figure 2, the amplitude of the P2 evoked by perception of the speech sounds was greater than the environmental sounds in the anterior ROIs but not in the posterior ROIs. Paired t-tests were conducted to inspect the interaction effects. These tests revealed that the mean voltage difference between the speech and environmental sounds was greatest in anterior regions (anterior – posterior: t_179_ = 9.7, p < 0.001) and in the middle of the scalp (middle – left: t_119_ = 4, p < .001, middle – right: t_119_ = 4.1, p < .001).

To determine if there were within-category perceptual differences driving the neural results, we conducted four separate ANOVAs. These ANOVAs compared the N1 and P2 amplitudes within category (dig vs. cut, car vs. frog). The factors were stimulus, anterior-posterior electrode position, and laterality. We found significant differences within category. The N1 evoked by the “dig” stimulus was more negative than that evoked by the “cut” stimulus in anterior regions. The N1 evoked by the frog stimulus was more negative than the car stimulus in anterior regions. The P2 was evoked by the car stimulus was greater than that evoked by the frog stimulus in anterior and middle regions (see Appendix).

#### 3.2.2 Imagery

Next, we tested whether there were categorical differences in the ERPs evoked by imagery of the speech sounds and nonspeech sounds. Grand average ERPs evoked by imagery of the speech and environmental sounds are displayed in Figure 3. We ran repeated measures ANOVAs with stimulus category, anterior-posterior electrode position, and laterality as factors and mean voltage across the LPC time window (350-500ms) as the dependent variable. This revealed main effects of stimulus category (F_1, 29_ = 8.770, p < .01) and anterior-posterior position (F_1, 29_ = 11.909, p < .01) and an interaction effect of anterior-posterior position and laterality (F_1.849, 53.609_ = 10.579, p < .001). As seen in Figure 3, imagery of environmental sounds elicited a greater LPC than speech sounds. There was no significant interaction between stimulus category and anterior-posterior position or stimulus category and laterality. However, paired t-tests were conducted contrasting laterality (left-middle, middle-right, and left-right) in the anterior and posterior regions to inspect their interaction effect. The amplitude of the LPC was significantly greater at the left (t_59_ = 3, p < .05) and middle (t_59_ = 4.5, p < .001) than the right electrodes in the posterior region. However, in the anterior region, the amplitude of the LPC was significantly greater at the right electrodes than the middle electrodes (t_59_ = 2.7, p < .05).

**Figure 3.**
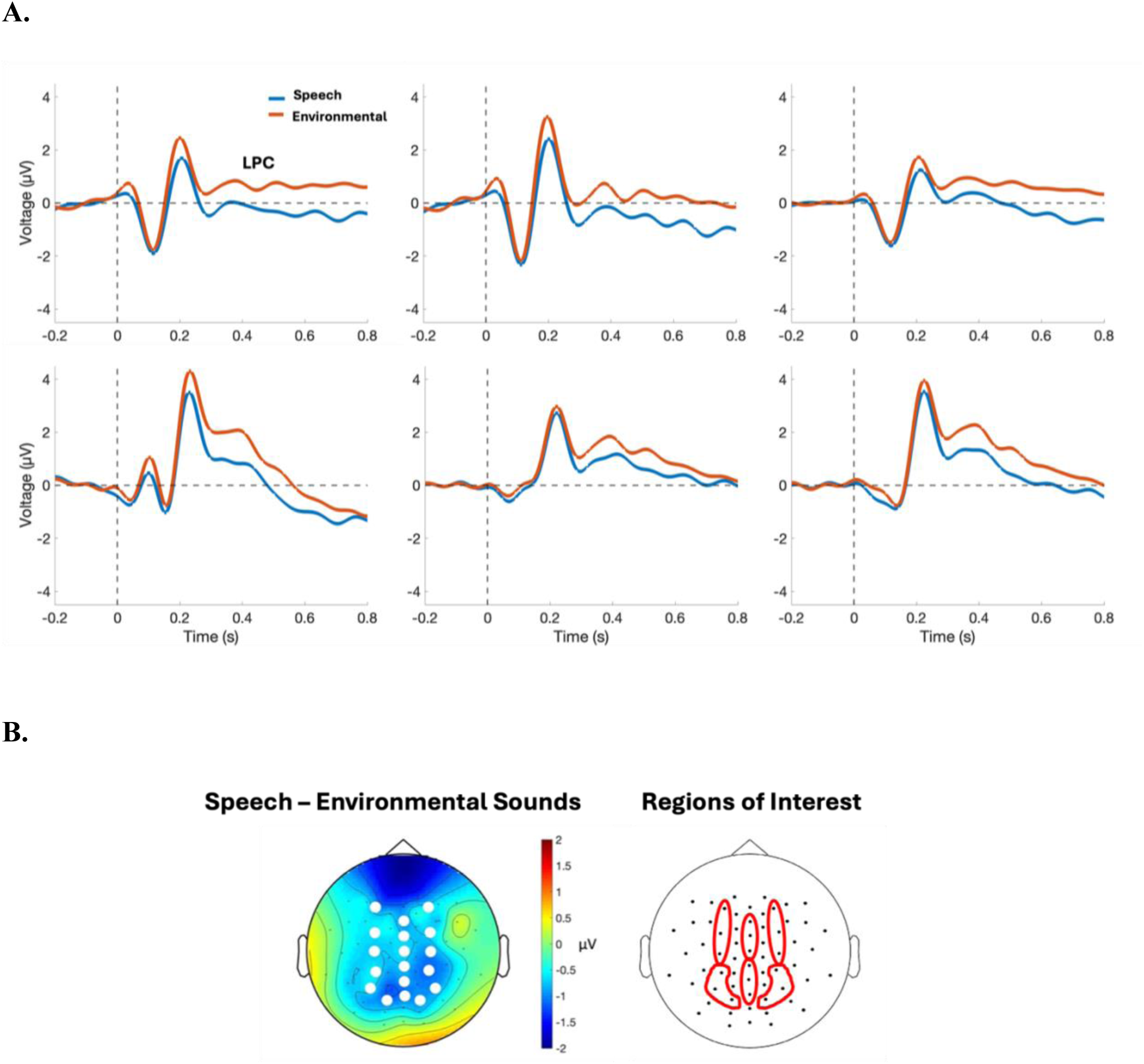
A. Grand average ERPs evoked by imagery of the speech and environmental sounds. ERPs are displayed from each ROI. B. The topographic plots represent the response to speech sounds minus the response to environmental sounds in the time window of interest (LPC). All 6 ROIs visually represented with red lines.

We investigated within-category differences in imagery responses in a similar way to perception responses. Four separate ANOVAs compared the N1 and P2 amplitudes within category (dig vs. cut, car vs. frog). The factors were stimulus, anterior-posterior electrode position, and laterality. We found no significant differences within category.

cvalidate the ERP results that showed differences between imagery of the speech sounds and the environmental sounds using an unbiased data-driven approach. The EEGNet algorithm (Lawhern et al., 2018) classified neural responses of speech sound imagery and environmental sound imagery with an average classification accuracy of 62%. To establish statistical significance, we employed a permutation approach where we randomly shuffled the condition labels and performed the classification. This null distribution yielded an accuracy of 50%, which was significantly lower than the real accuracy (p < .001). The spatio-temporal regions that contributed more weight for the classification between speech and nonspeech sounds were in posterior left and the anterior middle ROIs, specifically during the LPC window (Figure 4).

**Figure 4.**
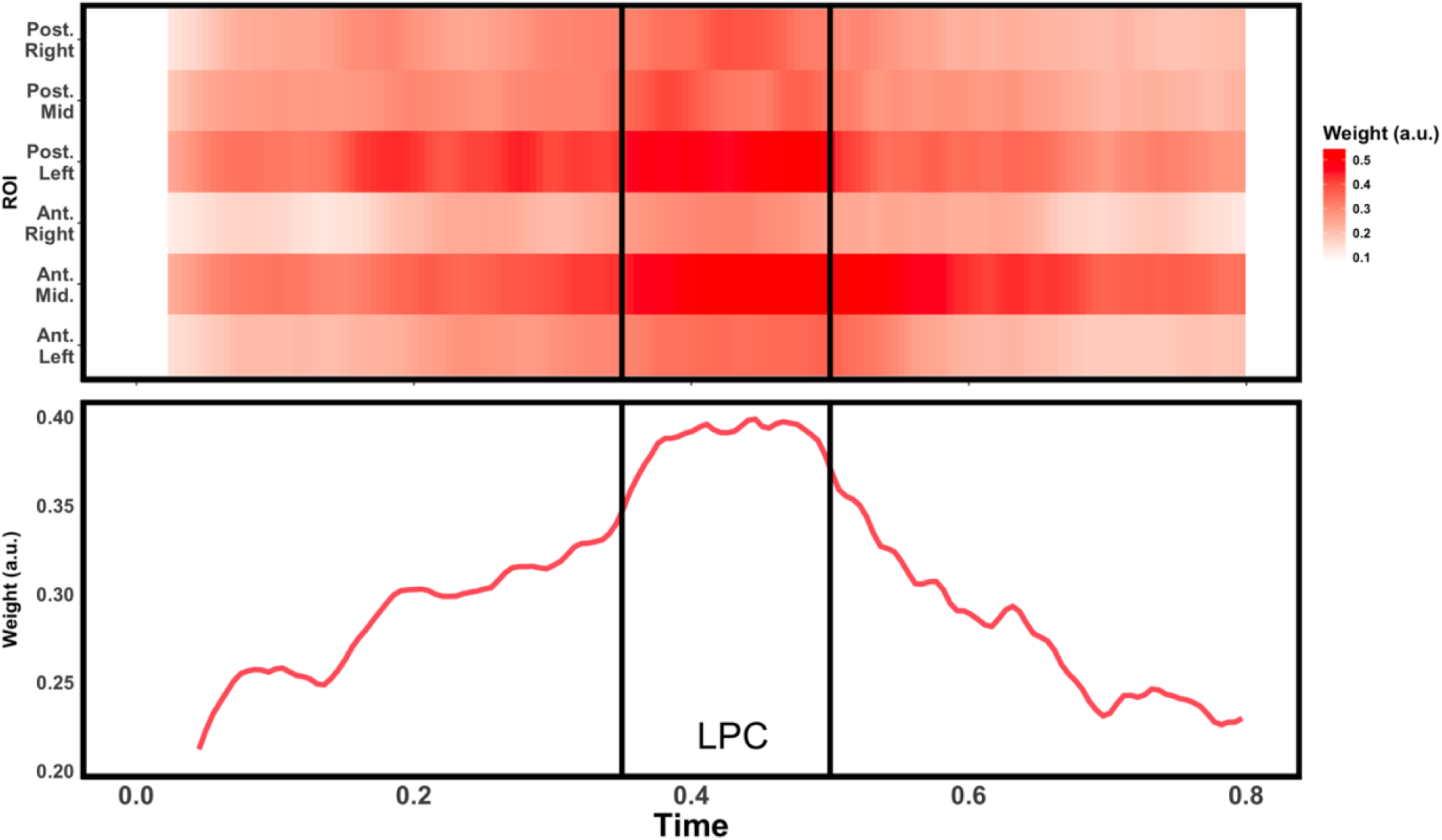
The top panel shows the average weights that the machine-learning algorithm assigned to electrodes within each ROI as a function of post-stimulus time. The bottom panel shows the weights averaged across ROIs. The vertical black lines delineate the period corresponding to the LPC window.

## 4 Discussion

The present study both confirms the presence of categorical processing in auditory perception and extends this processing to auditory imagery. We found early ERP component differences in perception of the speech as compared to environmental sounds and late differences in imagery. This later time window in imagery also provided salient information for a convolutional neural network classifying neural responses as imagery of speech or environmental sounds. These results contribute to our understanding of imagery — specifically auditory imagery — and its connection with perception.

### 4.1 Speech and Nonspeech as Perceptual Categories

The amplitude of the P2 ERP component was significantly greater for the perception of the speech sounds as compared to the environmental sounds across multiple ROIs. This supports previous research showing that human vocal sounds elicit a greater amplitude in early to mid-latency ERP components (Charest et al., 2009; Stavropoulos & Carver, 2016). This categorical difference could result from how the superior temporal sulcus has been shown to react with greater neuronal activity when processing human vocal sounds compared to environmental sounds (Belin et al., 2000). These results were further supported by the speech sounds eliciting a less negative amplitude N1 component than the environmental sounds. As in Charest et al. (2009), we suggest that this may be due to a lower power of human voice sounds at sound onset. Our results support the claim that human speech sounds are processed differently from environmental sounds in perception.

### 4.2 Neural Representation of Sound Categories in Imagery

The present study confirms that auditory imagery of sounds from different categories engages different neural processes. The amplitude of the LPC component was significantly greater for imagination of the environmental sounds as compared to imagination of the speech sounds across the central ROI. This supports our hypothesis that the LPC would reflect categorical differences in auditory imagery.

In Wu et al. (2011), the differences in the LPC evoked by imagery were in the same direction as the differences in the N1 evoked by perception. However, in the present study, although speech sounds elicited a greater amplitude P2 in perception, they elicited a lower amplitude LPC in imagery. One possible reason for this result could be that auditory imagery of

lexically meaningful speech sounds may not require as vivid a reproduction of the perceptual aspects of the stimulus. Previous research has shown that auditory imagery of music containing lyrics does not activate the primary auditory cortex, while auditory imagery of music without lyrics does activate the primary auditory cortex (Kraemer et al., 2005). We may see a similar effect for lexically meaningful speech, with lexical information being sufficient to produce an accurate auditory image without recruiting the primary auditory cortex. The time course of imagery differs from that of perception. Auditory perception relies on bottom-up activation of a pathway beginning in the temporal gyrus and proceeding to more frontal regions, while auditory imagery relies on top-down activation of a pathway beginning with the inferior frontal gyrus (IFG) and proceeding to the temporal gyrus (Ding et al., 2019). Imagery of lexically meaningful speech may not require reactivation of detailed sensory information, as we may rely on earlier lexical reconstruction in the IFG.

In contrast, the amplitude of the LPC may be modulated by the hierarchical structure of speech rather than lexical meaning. Previous research on categorical processing in auditory imagery has demonstrated that an ERP component similar to the LPC is suppressed in hierarchically complex stimuli, such as speech and music (Proverbio et al., 2023). Although Proverbio et al. (2023) suggested this component reflected a P2-like response, the amplitude was greater for emotional vocalizations than both music and lexically meaningful speech. It is not clear whether this categorical difference was due to the emotional valence of the vocal stimuli or the hierarchical complexity of the speech and musical stimuli. Further research is needed to determine whether late categorical differences in auditory imagery are due to lexical information, hierarchical structure, or emotional valence.

Another possibility is that the differences found in the LPC for imagery reflect the differences found in the N1 for perception. Properties that affect the amplitude of the N1 evoked by auditory perception — such as pitch and loudness — can affect the amplitude of the LPC (Wu et al., 2011). However, our stimuli were matched for pitch and loudness. This difference may reflect inherent differences in the acoustic structure of speech and environmental sounds, such as power at onset (Charest et al., 2009).

Further, the LPC was found to contain salient information for a convolutional neural network classifying neural responses as evoked by imagination of speech or environmental sounds. This result contributes to a growing literature on decoding imagined speech. Recent research has demonstrated the potential for neural speech prosthetics that decode imagined speech using ECoG (Proix et al., 2022). Imagined speech can be decoded from cortical areas outside the cortical language production network in patients with epilepsy, providing a foundation for brain-computer interfaces for individuals with lesions in this network due to stroke. While imagining oneself speaking recruits fronto-parietal sensorimotor regions, imagining hearing speech more strongly activates the inferior parietal cortex and intraparietal sulcus. Proix et al. (2022) interpreted this finding to suggest that imagined speaking involves a transformation from sensory to motor information, while imagined hearing relies more on memory retrieval of sensory information.

However, this research has been primarily conducted using invasive methods. The present study advances this work by using noninvasive electrophysiological recordings to examine the neural correlates of imagining different categories of sound. The present results demonstrate that EEG can differentiate between imagery of different categories of sound, suggesting the possibility of noninvasive decoding of auditory imagery.

### 4.3 Conclusion

The results of the present study suggest the presence of auditory category distinctions in both perception and imagery. Our results for perception contribute to existing evidence that auditory perception of human sounds functions via different neural processes from that of environmental sounds. More specifically, these results confirm previous research that has shown that the P2 ERP component elicited by perception of human vocal sounds is greater than that of nonhuman sounds (Charest et al., 2009; Stavropoulos & Carver, 2016).

In addition, our results for imagery expand upon previous research to suggest that auditory imagery of human sounds functions via different neural processes from that of nonhuman stimuli. Specifically, we showed that the nonspeech environmental sounds elicited a greater LPC component than the human speech sounds. Synthesis with previous research on categorical processing in auditory imagery can provide a more comprehensive explanation of this difference. For example, previous research has shown that spontaneous imagery of music that does not contain lyrics recruits the primary auditory cortex, while music containing lyrics does not (Kraemer et al., 2005). In addition, previous research has shown that human emotional vocalizations elicit a greater LPC (400-600 ms) than both lexically meaningful speech and musical stimuli (Proverbio et al., 2023). This suggests that the greater response to nonhuman stimuli in this study may be due to either the lexicality or the hierarchical structure of human speech stimuli. To test this, future research should be conducted comparing auditory imagery of lexically meaningful speech with lexically meaningless speech (ex. speech in an unfamiliar language) and nonspeech human vocal sounds. If suppression of the LPC is due to lexicality, auditory imagery of lexically meaningless speech and nonspeech vocal sounds would elicit a greater LPC than that of lexically meaningful speech. However, if this suppression is due to the hierarchical structure of speech, nonspeech vocal sounds would elicit a greater LPC than both lexically meaningful and meaningless speech.

Overall, the present study has demonstrated the presence of auditory category distinctions in both perception and imagery. Further research is needed to determine whether these differences are due to the lexicality of the stimuli, hierarchical structure of speech, or differential processing of the human voice. However, the present research contributes to a growing body of knowledge about imagery and its connection with perception.

## Data Code and Availability

Data and code available upon request.

## Author Contributions

**Julia M. Leeman:** Conceptualization, Methodology, Software, Formal Analysis, Investigation, Writing – Original Draft, Writing – Review & Editing, Visualization, Funding Acquisition; **Evan Hare:** Conceptualization, Methodology, Software, Investigation, Writing – Review & Editing, Visualization, Supervision, Project Administration; **Ricardo Morales-Torres:** Conceptualization, Methodology, Software, Formal Analysis, Writing – Review & Editing, Visualization, **Pooja Kabber:** Software, Formal Analysis, Investigation, **Joseph Zhang:** Software, Formal Analysis, Investigation; **Kristi Van Meter:** Software, Formal Analysis; **Tobias Overath:** Conceptualization, Methodology, Resources, Writing – Review and Editing, Supervision, Funding Acquisition

## Acknowledgements

We thank the Data+ program through the Rhodes Information Initiative at Duke University for facilities and partial funding.

## Appendix

### Validation of Stimuli

Analyses of variance (ANOVA) were conducted to determine whether reported difficulty of imagery varied by stimulus. The Type III analysis of variance using Satterthwaite’s method for difficulty of imagination revealed an effect of stimulus that trended toward significance (p = .06, F = 2.174). The contrasts did not reveal a significant difference in difficulty of imagination between any two stimuli after adjusting the p-values to control for the false discovery rate; however, the chicken and screenshot stimuli had the highest reported difficulty (see Supplementary Figure 1). For this reason, the chicken and screenshot stimuli were removed from analysis. There was no significant effect of vividness.

**Supplementary Figure 1.**
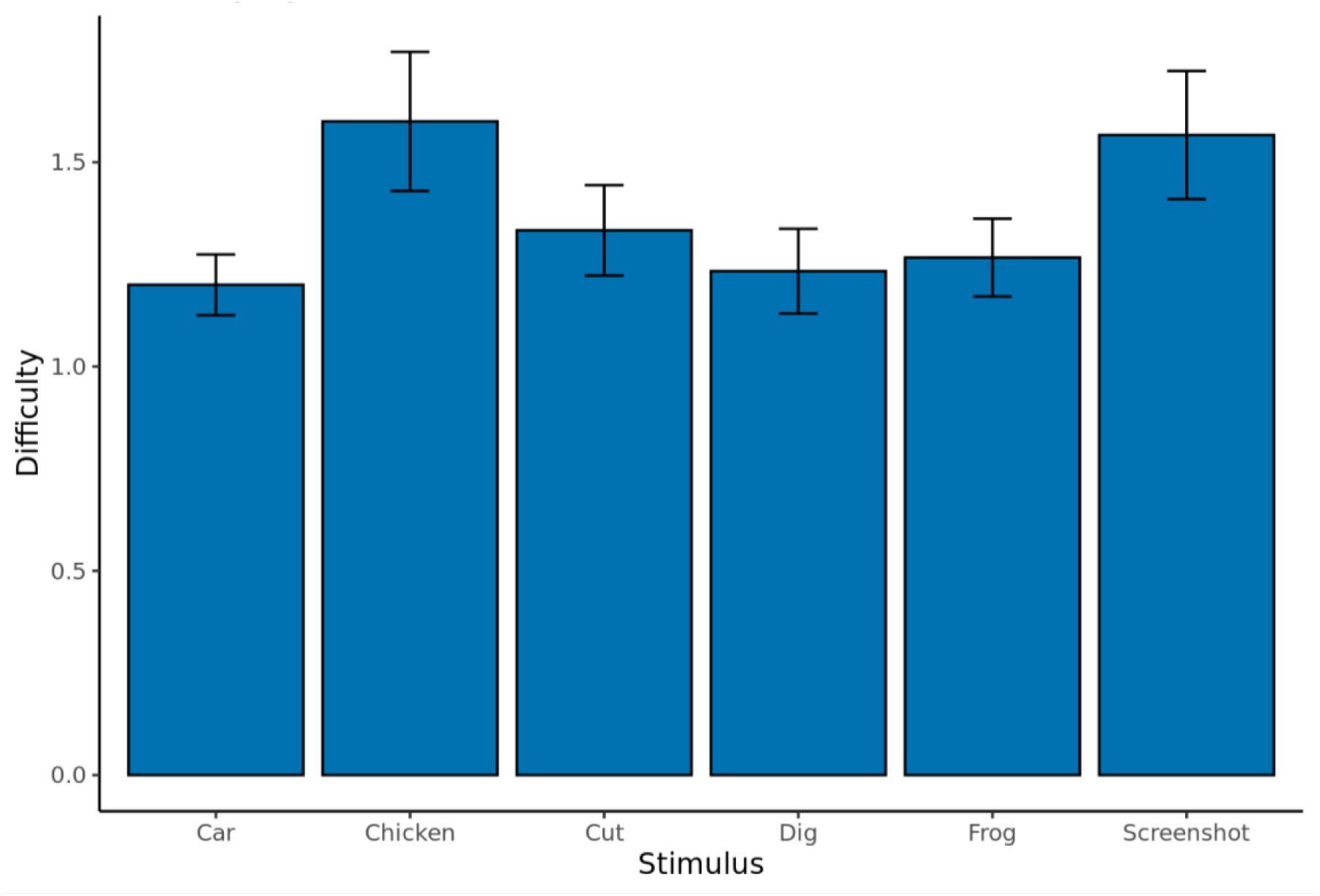
Reported difficulty of imagination by stimulus. Chicken and Screenshot stimuli were removed from analysis due to greater difficulty of imagination.

### Within-Category ERP Analysis

We found a main effect of stimulus (F_1, 29_ = 5.146, p < .05) and an interaction effect of stimulus and anterior-posterior electrode position (F_1, 29_ = 4.599, p < .05) when comparing the amplitude of the N1 evoked by perception of the two speech stimuli. A paired t-test revealed that the mean voltage difference between the cut and dig stimuli was greatest in anterior regions (anterior - posterior: t_89_ = 3.6, p < .001), with dig stimuli having a more negative N1 than cut stimuli in these regions. We found an interaction effect of stimulus and anterior-posterior electrode position (F_1, 29_ = 14.670, p < .001) when comparing the amplitude of the N1 evoked by perception of the two nonspeech stimuli. A paired t-test revealed that the mean voltage difference between the car and frog stimuli was greatest in anterior regions (anterior - posterior: t_89_ = 6.2, p < .001), with frog stimuli having a more negative N1 than car stimuli in these regions.

Additionally, we found interaction effects of stimulus and anterior-posterior electrode position (F_1, 29_ = 12.598, p < .01) and of stimulus and laterality (F_2, 58_ = 4.243, p < .05) when comparing the amplitude of the P2 evoked by perception of the two nonspeech stimuli. Paired t-tests were conducted to inspect the interaction effects. These tests revealed that the mean voltage difference between the car and frog stimuli was greatest in anterior regions (anterior - posterior: t_89_ = 5.8, p < .001) and in the middle of the scalp (middle - left: t_59_ = 3, p < .01, middle - right: t_59_ = 2.9, p < .01), with car stimuli having a greater amplitude P2 than frog stimuli in these regions.

## References

1. Ancona, M., Ceolini, E., Öztireli, C., & Gross, M. (2018). Towards better understanding of gradient-based attribution methods for Deep Neural Networks (No. arXiv:1711.06104). arXiv. 10.48550/arXiv.1711.06104

2. Belin, P., Zatorre, R. J., Lafaille, P., Ahad, P., & Pike, B. (2000). Voice-selective areas in human auditory cortex. Nature, 403(6767), 309–312. 10.1038/35002078

3. Brainard, D. H. (1997). The Psychophysics Toolbox. Spatial Vision, 10(4), 433–436.

4. Charest, I., Pernet, C. R., Rousselet, G. A., Quiñones, I., Latinus, M., Fillion-Bilodeau, S., Chartrand, J.-P., & Belin, P. (2009). Electrophysiological evidence for an early processing of human voices. BMC Neuroscience, 10(1), 127. 10.1186/1471-2202-10-127

5. Delorme, A., & Makeig, S. (2004). EEGLAB: An open source toolbox for analysis of single-trial EEG dynamics including independent component analysis. Journal of Neuroscience Methods, 134(1), 9–21. 10.1016/j.jneumeth.2003.10.009

6. Dijkstra, N., Bosch, S. E., & Gerven, M. A. J. van. (2017). Vividness of Visual Imagery Depends on the Neural Overlap with Perception in Visual Areas. Journal of Neuroscience, 37(5), 1367–1373. 10.1523/JNEUROSCI.3022-16.2016

7. Dijkstra, N., Bosch, S. E., & van Gerven, M. A. J. (2019). Shared Neural Mechanisms of Visual Perception and Imagery. Trends in Cognitive Sciences, 23(5), 423–434. 10.1016/j.tics.2019.02.004

8. Ding, Y., Zhang, Y., Zhou, W., Ling, Z., Huang, J., Hong, B., & Wang, X. (2019). Neural Correlates of Music Listening and Recall in the Human Brain. Journal of Neuroscience, 39(41), 8112–8123. 10.1523/JNEUROSCI.1468-18.2019

9. Gelding, R. W., Thompson, W. F., & Johnson, B. W. (2019). Musical imagery depends upon coordination of auditory and sensorimotor brain activity. Scientific Reports, 9(1), 16823. 10.1038/s41598-019-53260-9

10. Halpern, A. R. (2015). Differences in auditory imagery self-report predict neural and behavioral outcomes. *Psychomusicology: Music*, Mind, and Brain, 25(1), 37–47. 10.1037/pmu0000081

11. Hubbard, T. L. (2010). Auditory imagery: Empirical findings. Psychological Bulletin, 136(2), 302–329. 10.1037/a0018436

12. Kraemer, D. J. M., Macrae, C. N., Green, A. E., & Kelley, W. M. (2005). Sound of silence activates auditory cortex. Nature, 434(7030), 158–158. 10.1038/434158a

13. Lawhern, V. J., Solon, A. J., Waytowich, N. R., Gordon, S. M., Hung, C. P., & Lance, B. J. (2018). EEGNet: A compact convolutional neural network for EEG-based brain-computer interfaces. Journal of Neural Engineering, 15(5), 056013. 10.1088/1741-2552/aace8c

14. Levy, D. A., Granot, R., & Bentin, S. (2001). Processing specificity for human voice stimuli: Electrophysiological evidence. Neuroreport, 12(12), 2653–2657. 10.1097/00001756-200108280-00013

15. McNorgan, C. (2012). A meta-analytic review of multisensory imagery identifies the neural correlates of modality-specific and modality-general imagery. Frontiers in Human Neuroscience, 6. 10.3389/fnhum.2012.00285

16. Oostenveld, R., Fries, P., Maris, E., & Schoffelen, J.-M. (2011). FieldTrip: Open source software for advanced analysis of MEG, EEG, and invasive electrophysiological data. Computational Intelligence and Neuroscience, 2011, 156869. 10.1155/2011/156869

17. Proix, T., Delgado Saa, J., Christen, A., Martin, S., Pasley, B. N., Knight, R. T., Tian, X., Poeppel, D., Doyle, W. K., Devinsky, O., Arnal, L. H., Mégevand, P., & Giraud, A.-L. (2022). Imagined speech can be decoded from low- and cross-frequency intracranial EEG features. Nature Communications, 13(1), 48. 10.1038/s41467-021-27725-3

18. Proverbio, A. M., Tacchini, M., & Jiang, K. (2023). What do you have in mind? ERP markers of visual and auditory imagery. Brain and Cognition, 166, 105954. 10.1016/j.bandc.2023.105954

19. Regev, M., Halpern, A., Owen, A., Patel, A., & Zatorre, R. (2021). Mapping Specific Mental Content during Musical Imagery. Cerebral Cortex, 31(8), 3622–3640.

20. Stavropoulos, K. K.-M., & Carver, L. J. (2016). Neural Correlates of Attention to Human-Made Sounds: An ERP Study. PLOS ONE, 11(10), e0165745. 10.1371/journal.pone.0165745

21. The MathWorks Inc. (2018). MATLAB version: 9.5.0 (R2018b). Natick, Massachusetts: The MathWorks Inc. https://www.mathworks.com

22. The MathWorks Inc. (2021). MATLAB version: 9.11.0 (R2021b). Natick, Massachusetts: The MathWorks Inc. https://www.mathworks.com

23. Tian, X., Ding, N., Teng, X., Bai, F., & Poeppel, D. (2018). Imagined speech influences perceived loudness of sound. Nature Human Behaviour, 2(3), 225–234. 10.1038/s41562-018-0305-8

24. Tsunada, J., & Cohen, Y. E. (2014). Neural mechanisms of auditory categorization: From across brain areas to within local microcircuits. Frontiers in Neuroscience, 8. 10.3389/fnins.2014.00161

25. Tużnik, P., Augustynowicz, P., & Francuz, P. (2018). Electrophysiological correlates of timbre imagery and perception. International Journal of Psychophysiology, 129, 9–17. 10.1016/j.ijpsycho.2018.05.004

26. Wu, J., Yu, Z., Mai, X., Wei, J., & Luo, Y. (2011). Pitch and loudness information encoded in auditory imagery as revealed by event-related potentials. Psychophysiology, 48(3), 415–419. 10.1111/j.1469-8986.2010.01070.x

27. Zatorre, R. J., Halpern, A. R., & Bouffard, M. (2010). Mental Reversal of Imagined Melodies: A Role for the Posterior Parietal Cortex. Journal of Cognitive Neuroscience, 22(4), 775– 789. 10.1162/jocn.2009.21239

